# Upregulated Fcrl5 disrupts B cell anergy and causes autoimmunity

**DOI:** 10.1101/2023.07.20.549843

**Authors:** Chisato Ono, Shinya Tanaka, Keiko Myouzen, Takeshi Iwasaki, Mahoko Ueda, Yoshinao Oda, Kazuhiko Yamamoto, Yuta Kochi, Yoshihiro Baba

## Abstract

B cell anergy plays a critical role in maintaining self-tolerance by inhibiting autoreactive B cell activation to prevent autoimmune diseases. Here, we demonstrated that Fc receptor-like 5 (Fcrl5) upregulation contributes to autoimmune disease pathogenesis by disrupting B cell anergy. Fcrl5—a gene whose homologs are associated with human autoimmune diseases—is highly expressed in age/autoimmunity-associated B cells (ABCs), an autoreactive B cell subset. By generating B cell-specific Fcrl5 transgenic mice, we demonstrated that Fcrl5 overexpression in B cells caused systemic autoimmunity with age. Furthermore, Fcrl5 upregulation in B cells exacerbated the systemic lupus erythematosus-like disease model and increased the levels of ABCs and activated T cells. Mechanistically, an increase in Fcrl5 expression broke B cell anergy, activating autoreactive B cells in the presence of a self-antigen. Fcrl5 facilitated toll-like receptor signaling, reactivating anergic B cells. Thus, Fcrl5 is a potential regulator of B cell-mediated autoimmunity by regulating B cell anergy. This study provides important insights into the role of Fcrl5 in breaking B cell anergy and its effect on the pathogenesis of autoimmune diseases.

## Introduction

B cells play a crucial role in promoting various autoimmune diseases by secreting autoantibodies, presenting autoantigens to T cells, and producing inflammatory cytokines (Hao et al., 2011; Rubtsov et al., 2015). These pathogenic B cells are considered autoreactive and express B cell antigen receptors (BCRs) with self-specificity, which is common in the large proportion of newly formed B cells (Cambier et al., 2007; Nemazee, 2017; Wardemann et al., 2003). These B cells should be censored to avoid or ameliorate autoimmune diseases caused by multiple immune tolerance systems, including clonal deletion, BCR editing, and autoreactive B cell anergy. B cell anergy is a state of functional inactivation or unresponsiveness in B cells, which is the primary mechanism operating in the periphery and is induced by chronic exposure to self-antigens in autoreactive B cells (Ferry et al., 2006; Goodnow et al., 1988). The anergic state is reversible and evaded by stimuli other than BCRs, including signals from toll-like receptors (TLRs), resulting in the accumulation of autoreactive B cells and an increased risk of developing autoimmune diseases (Theofilopoulos et al., 2017; Rawlings et al., 2017). One factor in the pathogenesis of systemic autoimmune lupus is the poor clearance of dead cell debris, releasing RNA and DNA, which activate endosomal TLR7 and TLR9 signaling, respectively (Tsokos et al., 2016).

Memory B and plasma cells are pathogenic in autoimmune diseases; however, a new subset of cells—age/autoimmunity-associated B cells (ABCs)—has recently been suggested to contribute to autoimmunity (Li et al., 2023; Mouat et al., 2022). ABCs were initially identified as a cell population that increased with age; however, they expanded in mouse autoimmune models regardless of age. In addition to conventional B cell markers, ABCs exhibit a characteristic phenotype in which surface CD21 and CD23 are absent, whereas the myeloid markers CD11b and CD11c and the transcription factor T-bet are often present (Hao et al., 2011; Rubtsov et al., 2011). ABCs are phenotypically heterogeneous and exhibit the properties of antigen-exposed cells and memory B cells. ABCs are often variedly defined depending on the combination of these markers. Human ABC-like B cells (including CD27^−^IgD^−^ and CD11c^+^CD21^lo^) are more abundant in various human autoimmune diseases, such as systemic lupus erythematosus (SLE), and rheumatoid arthritis (RA) (Li et al., 2023; Mouat et al., 2022). However, molecules expressed in ABCs that are involved in autoimmune diseases are unknown.

Genome-wide association studies of autoimmune diseases have identified many loci associated with disease risks, shedding light on the development of each disease. Among them, single nucleotide polymorphisms (SNPs) in genes such as human Fc receptor-like 3 (*hFcrl3*) or *hFcrl5* are associated with many autoimmune diseases such as RA (Kochi et al., 2005), SLE (Gibson et al., 2009; Mao et al., 2010), multiple sclerosis (Yuan et al., 2016), Graves’ disease (Simmonds et al., 2006, 2010), and Hashimoto’s disease (Kalantar et al., 2020). Notably, SNPs in the promoter region of hFcrl3 upregulate gene expression, which may be a risk factor for autoimmune diseases (Kochi et al., 2005). In humans, eight hFcrls—hFcrl1-6, hFcrlA, and hFcrlB—exist, whereas in mice, only six family members—Fcrl1, Fcrl5, Fcrl6, Fcrls, Fcrla, and Fcrlb—are present (Maltais et al., 2006). hFcrl3 and hFcrl5 share a significant degree of sequence and domain homology with mouse Fcrl5. They contain an immunoreceptor tyrosine-based activation motif (ITAM) and an immunoreceptor tyrosine-based inhibition motif (ITIM) in their intracellular regions (Li et al., 2014), which may play opposing roles in regulating signaling pathways downstream of BCRs. Mouse Fcrl5 is expressed in B-1 cells, marginal zone (MZ) B cells, and atypical memory B cells, and depending on the cells in which it is expressed, it may play an activating or inhibitory role (Zhu et al., 2013; Kim et al., 2019). However, the role of Fcrl5 expression in B cells in the pathogenesis of autoimmune disease remains unknown.

Here, we observed the high expression of Fcrl5 in the ABCs of aged mice. We found that B cell-specific Fcrl5 transgenic (Tg) mice overexpressing this gene developed systemic autoimmunity with age and exacerbated SLE-like disease model. Mechanistically, Fcrl5 overexpression disrupted B cell anergy and facilitated TLR signaling, possibly contributing to autoimmunity.

## Results

### Identification of differentially expressed genes in ABCs

To explore gene expression in ABCs, we performed RNA-sequencing (RNA-seq) on the sorted splenic ABCs and follicular (FO) B cells of aged mice and FO B cells of young mice (Fig. S1 A). Here, we defined ABCs as CD19^+^B220^hi^CD43^−^CD23^−^CD21^lo^CD11b^+^CD11c^+^ B cells to exclude other subsets such as B1, transitional, and memory B cells (Riedel et al., 2020). A comparison of RNA-seq data between ABCs and FO B cells in aged mice showed 152 upregulated and 40 downregulated genes in ABCs, with a greater than fivefold expression difference (Fig. 1 A and B). FO B cells from young and aged mice showed similar gene expression patterns (Fig. 1 B and Fig. S1 B). Among the ABC upregulated genes, we focused on Fcrl5, an ortholog of hFcrl3 whose upregulated expression is closely associated with autoimmune disease development (Li et al., 2014; Kochi et al., 2005). The expression quantitative trait loci (eQTL) catalog in ImmuNexUT—a comprehensive database consisting of RNA-seq data and the eQTL database of immune cells from patients with immune-mediated diseases—revealed that the effects of eQTL on *hFcrl3* and *hFcrl5* expression in all cell subsets were observed in autoimmune disease specimens. It also revealed that *hFcrl3/5* transcripts were higher in the CD27^−^IgD^−^DN B cell subset (human ABC-like cells) (Fig. S1, C and D). The RNA-seq and quantitative polymerase chain reaction (qPCR) analyses revealed that among the Fcrl family members, Fcrl5 was the only upregulated gene in ABCs than in FO B cells (Fig. 1, C and D). Flow cytometry confirmed that Fcrl5 was expressed at low levels in FO B cells regardless of age but was highly expressed in ABCs (Fig. 1 E). As previously reported, Fcrl5 was highly expressed in MZ B, germinal center (GC) B, and plasma cells and was weakly expressed in other B cell subsets in the spleens of young and aged mice (Fig. S1 E). Moreover, Fcrl5 expression decreased slightly in B1b and B2 cells in the peritoneal cavity of aged mice compared with that in young mice (Fig. S1 F). Thus, ABCs, potential drivers of autoimmune diseases, show high Fcrl5 expression.

**Figure 1.**
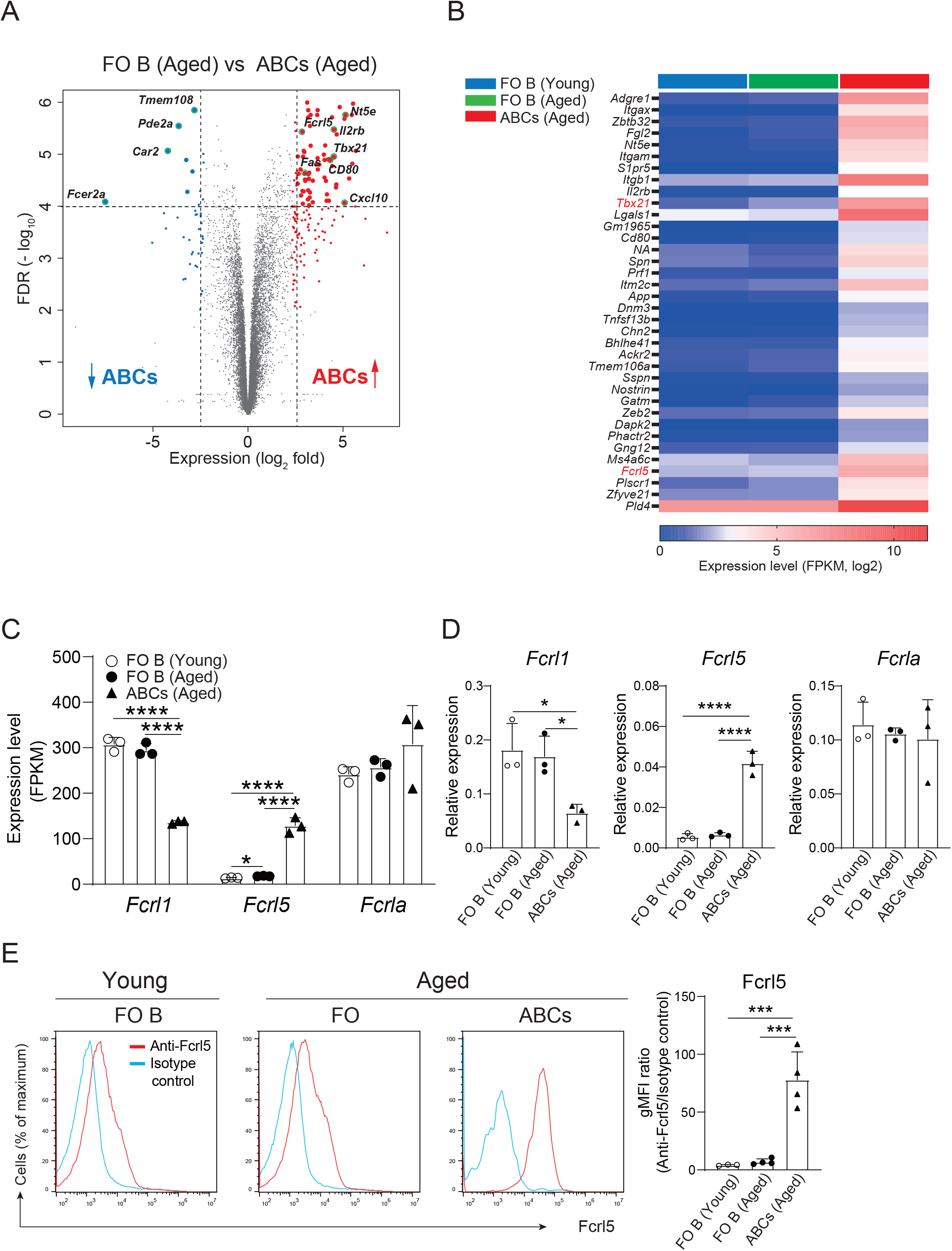
Identification of differentially expressed genes in ABCs. (A) Differential gene expression in FO B (CD19^+^B220^hi^CD43^−^CD23^+^CD21^lo^) and ABCs (CD19^+^B220^hi^CD43^−^CD23^−^CD21^lo^CD11b^+^CD11c^+^) from aged WT mice, presented as a volcano plot showing genes expressed in ABCs relative to their expression in FO B cells (n=3 mice each), plotted against log_2_ fold change >2.3 and FDR<0.01. (B) Selected gene expression of the highly expressed gene in ABCs compared to FO B cells sorted from young and aged WT mice (n=3 mice each) with the parameters; log_2_ fold change >2.5, FDR<0.01 and adjusted P value <0.00002. (C) Relative expression (FPKM) of *Fcrl1, Fcrl5,* and *Fcrla* mRNA in sorted FO B cells and ABCs from young or aged WT mice (n=3 mice each). FPKM, fragments per kilobase of transcript per million reads mapped. (D) Relative expression of *Fcrl1, Fcrl5,* and *Fcrla* analyzed by qPCR in sorted FO B cells and ABCs from young or aged WT mice (n=3 mice each). (E) Representative histograms (left) and gMFIs (right) of Fcrl5 expression on FO B cells and ABCs from young (n=3) or aged (n=4) WT mice. gMFI, geometric mean fluorescence intensity. Data are representative of three independent experiments. Statistical data are shown as mean values with s.d., and data were analyzed by one-way ANOVA with Tukey’s multiple comparisons test. *P<0.05, ***P<0.001, ****P<0.0001.

### B cell-specific Fcrl5 overexpression in mice develops autoimmune diseases with age

Upregulated hFcrl3 expression is associated with an increased risk of autoimmune diseases (Kochi et al., 2005). To elucidate the Fcrl5 upregulation effect on B cells, we generated B cell-specific Fcrl5 Tg mice (Fig. S2 A). We confirmed high expression of Fcrl5 in B cell lineages of the bone marrow (BM), peritoneal cavities (PeC), payer’s patches (PPs), and spleens of Fcrl5 Tg mice (Fig. S2 B). These mice were viable, fertile, born at the expected Mendelian frequencies, and exhibited normal B cell development in the BM, PeC, PPs, and spleens at a young age (Fig. S2 C). Serum immunoglobulin (Ig)M titers were higher in Fcrl5 Tg mice than in wild-type (WT) mice (Fig. S3 A).

Aging contributes to autoimmune disease development (de Mol et al., 2021). To determine whether Fcrl5 Tg mice showed autoimmune symptoms, we examined cellular and systemic inflammation-related abnormalities in aged mice (50–100-week-old). The aged Fcrl5 Tg mice showed higher IgM and IgA serum levels and lower IgG3 levels than those shown by age-matched WT mice (Fig. S3 A). B cell development in the spleen of aged Fcrl5 Tg mice was normal (Fig. S3 B). Compared with age-matched control WT mice, aged Fcrl5 Tg mice showed increased antinuclear antibodies (ANAs) production (Fig. 2 A) and serum titers of IgG against double-stranded DNA (dsDNA) (Fig. 2 B). Furthermore, hematoxylin and eosin staining revealed lymphocytic infiltrates in the liver, lung, and kidney with age. These infiltrates were more severe in Fcrl5 Tg mice (Fig. 2 C). These findings suggest that upregulated Fcrl5 expression in B cells disrupts the B cell-intrinsic program for immune tolerance.

**Figure 2.**
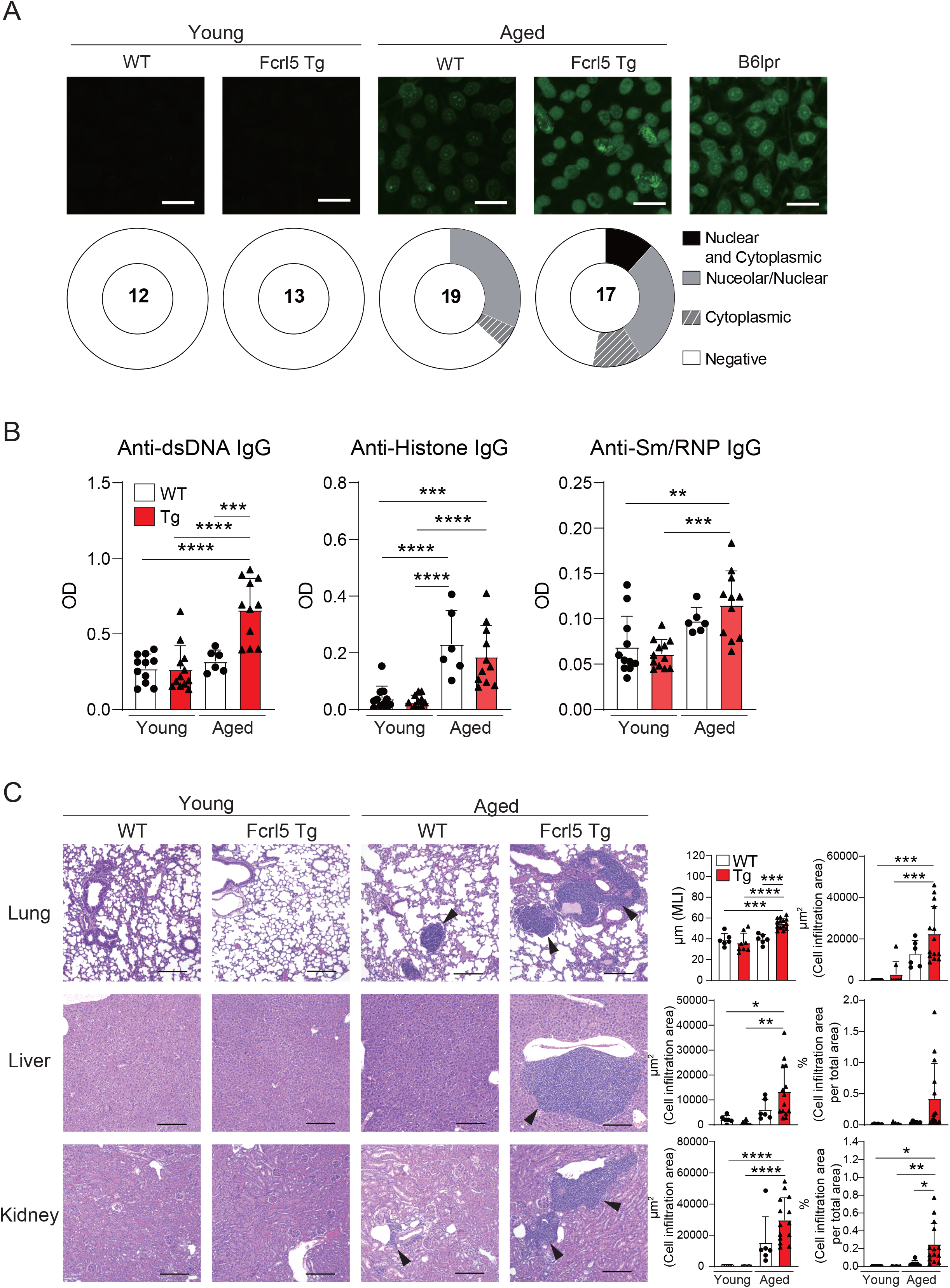
Aged Fcrl5 Tg mice develop autoimmune disease. (A) Representative images of ANA staining obtained with serum from young WT (n=12), young Fcrl5 Tg (n=13), aged WT (n=19), and aged Fcrl5 Tg (n=17) mice, detected by immunofluorescence assay using HEp-2 cells. Scale Bars, 50 μm. Data are pooled from three independent experiments. (B) Autoantibody production against dsDNA, histone, and Sm/RNP (ribonucleoprotein) in serum isolated from young WT (n=11), young Fcrl5 Tg (n=12), aged WT (n=6), and aged Fcrl5 Tg (n=11) mice, assayed by ELISA. OD, optical density. Data are pooled from three independent experiments. (C) Left, representative H&E-stained histological lung, liver, and kidney images of young WT (n=6), young Fcrl5 Tg (n=8), aged WT (n=6), and aged Fcrl5 Tg (n=14) mice. Arrowheads indicate areas of cell infiltration. Scale bars, 200 μm. Data are representative of three independent experiments. Right, quantitated cell infiltration (cell infiltration area per total area) in the lung, liver, and kidney and mean linear intercept (MLI) in the lung of young WT (n=6), young Fcrl5 Tg (n=8), aged WT (n=6), and aged Fcrl5 Tg (n=14) mice. Data are pooled from three independent experiments. Statistical data are shown as mean values with s.d., and data were analyzed by one-way ANOVA with Tukey’s multiple comparisons test. *P<0.05, **P<0.01, ***P<0.001, ****P<0.0001.

### Upregulation of Fcrl5 in B cells promotes autoimmune diseases

To investigate whether upregulated Fcrl5 expression affects autoimmune disease progression, we used the TLR7 agonist imiquimod-induced SLE-like model. We exposed 8–12-week-old WT and Fcrl5 Tg mice to imiquimod every 2 days for 10 weeks (Fig. 3 A). In imiquimod-treated Fcrl5 Tg mice, ANAs production increased compared with that in WT mice (Fig. 3 B). This increase was observed in the second week of induction (Fig. S4 A). The production of IgG autoantibodies against dsDNA and Sm/RNP increased in Fcrl5 Tg mice after imiquimod treatment (Fig. 3 C). H&E staining showed that imiquimod-treated Fcrl5 Tg mice showed extensive lymphocyte aggregation in the kidneys (Fig. 3 D). To gain insight into the effect of Fcrl5 expression in B cells on pathogenesis, we examined the cell activation status in the spleens of imiquimod-treated WT and Fcrl5 Tg mice. The number of CD19^+^ B cells, CD4^+^, and CD8^+^ T cells was increased in WT mice by imiquimod treatment (Fig. 4 A). The Fcrl5 Tg mice showed more progressive splenomegaly (data not shown) and increased numbers of CD19^+^ B cells and CD4^+^ T cells than WT mice (Fig. 4 A). Imiquimod treatment increased the number of CD23^−^CD21^lo^, GC B cells, and ABCs in WT mice, and this increase was even greater in Fcrl5 Tg mice (Fig. 4, B and C). The levels of B cell activation markers such as CD69, CD80, CD86, major histocompatibility complex (MHC) class Ⅱ, and programmed cell death ligand 1 (PD-L1) increased similarly in these mice after imiquimod treatment (Fig. S4 B). Additionally, activated CD4^+^ T, T follicular helper (Tfh), regulatory T (Treg), and interferon (IFN)-γ/interleukin (IL)-2-producing Th1 cells were augmented more in Fcrl5 Tg mice than in WT mice (Fig. 4 D), suggesting that Fcrl5 upregulation in B cells altered T cell activation under autoimmune conditions. These results indicate that upregulated Fcrl5 expression exacerbates the pathogenesis of SLE-like autoimmune diseases. Notably, we observed that imiquimod-induced lymphocyte activation and the increase in ABC levels were BCR-dependent, as assessed in MD4 Tg mice that expressed fixed membrane IgM specifically recognizing hen egg lysozyme (HEL) (Fig. S5, A-C). This suggests that TLR7-stimulated immune responses in various cells—including self-antigen-recognizing B cells—may play a pivotal role in the pathogenesis of this model and that Fcrl5 upregulation may potentially augment this response.

**Figure 3.**
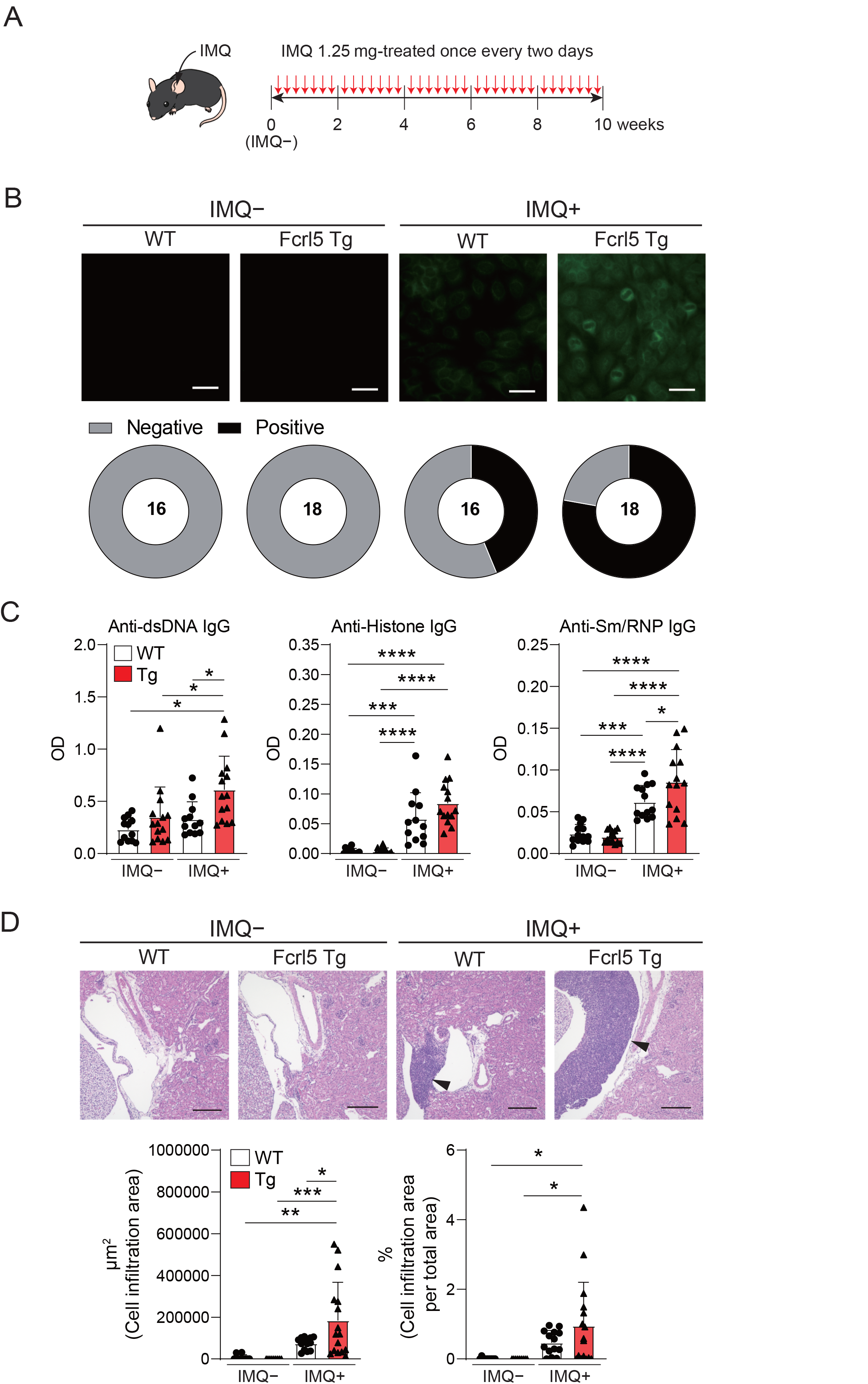
Upregulation of Fcrl5 in B cells exacerbates SLE-like disease model. (A) A scheme for induction of an imiquimod (IMQ)-induced SLE-like model. (B) Representative images of ANA staining obtained with serum from imiquimod-untreated (IMQ–) or -treated (IMQ+) WT (n=16) and Fcrl5 Tg (n=18) mice, detected by immunofluorescence assay using HEp-2 cells. Scale Bars, 50 μm. Data are pooled from three independent experiments. (C) Autoantibody production against dsDNA, histone, and Sm/RNP (ribonucleoprotein) in serum isolated from imiquimod-untreated (IMQ–) or -treated (IMQ+) WT (n=12) and Fcrl5 Tg (n=14) mice, assayed by ELISA. OD, optical density. Data are pooled from three independent experiments. (D) Representative H&E-stained histological kidney images and quantitated cell infiltration (cell infiltration area per total area) in the kidney of imiquimod-untreated (IMQ–) WT (n=9) and Fcrl5 Tg (n=9) or -treated (IMQ+) WT (n=14) and Fcrl5 Tg (n=16) mice. Arrowheads indicate areas of cell infiltration. Scale bars, 200 μm. Data are pooled from three independent experiments. Statistical data are shown as mean values with s.d., and data were analyzed by one-way ANOVA with Tukey’s multiple comparisons test. *P<0.05, **P<0.01, ***P<0.001, ****P<0.0001.

**Figure 4.**
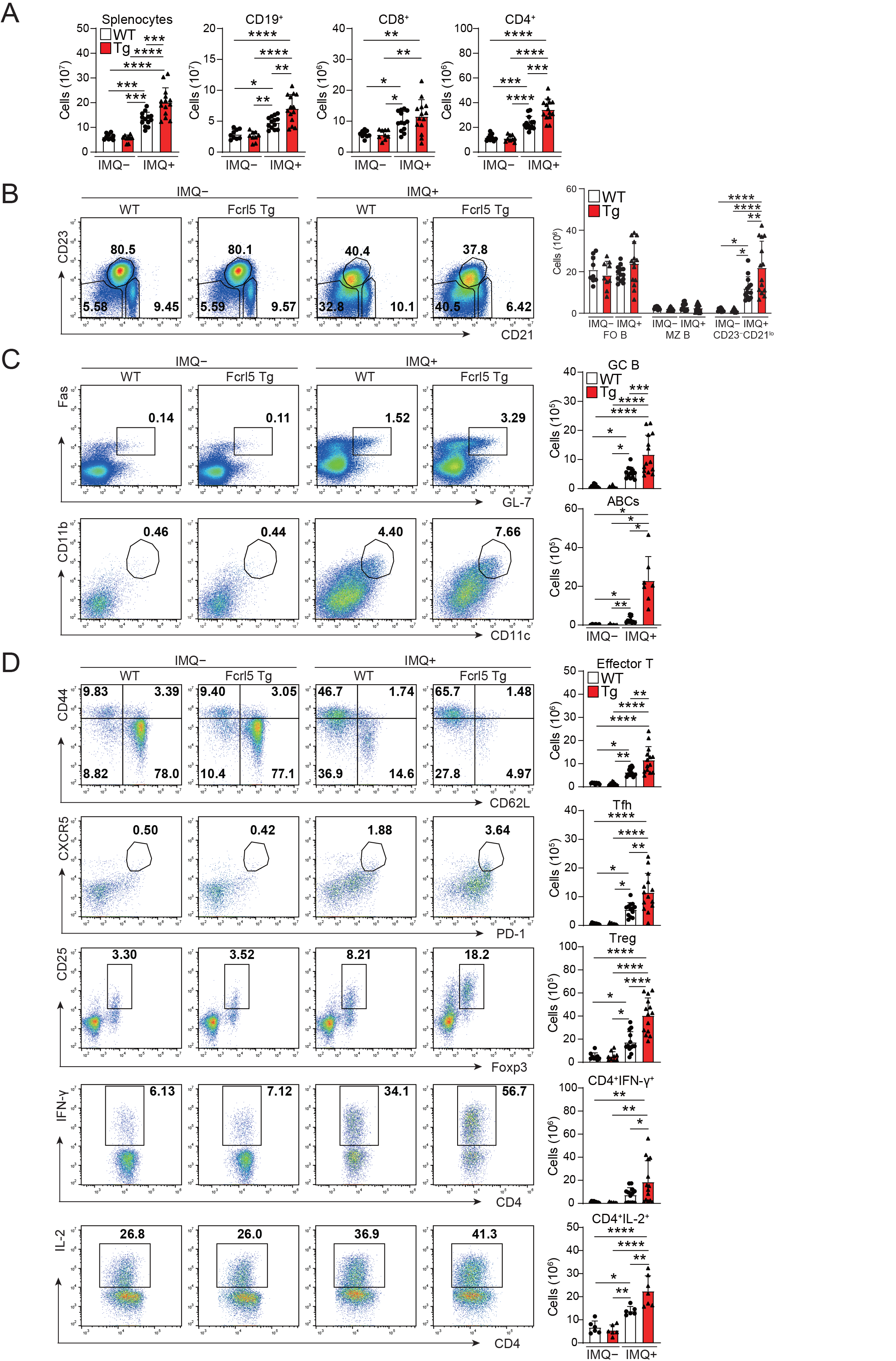
Enhanced activation of lymphocytes in imiquimod-treated Fcrl5 Tg mice. (A) Absolute number of splenocytes, CD19^+^, CD8^+^, and CD4^+^ in spleen from imiquimod-untreated (IMQ–) WT (n=9) and Fcrl5 Tg (n=9) or -treated (IMQ+) WT (n=13) and Fcrl5 Tg (n=14) mice. Data are pooled from three independent experiments. (B) Representative flow cytometry plots and absolute numbers of CD23^−^CD21^lo^ (CD19^+^CD93^−^CD23^−^CD21^lo^), FO B (CD19^+^CD93^−^ CD23^+^CD21^lo^), and MZ B (CD19^+^CD93^−^CD23^−^CD21^hi^) cells from imiquimod-untreated (IMQ–) WT (n=9) and Fcrl5 Tg (n=9) or -treated (IMQ+) WT (n=13) and Fcrl5 Tg (n=14) mice. Data are pooled from three independent experiments. (C) Top, representative flow cytometry plots, frequencies, and absolute numbers of GC B (B220^+^Fas^+^GL7^+^) cells from imiquimod-untreated (IMQ–) WT (n=9) and Fcrl5 Tg (n=9) or -treated (IMQ+) WT (n=13) and Fcrl5 Tg (n=14) mice. Bottom, representative flow cytometry plots and absolute numbers of ABCs (CD19^+^B220^hi^CD23^−^ CD21^lo^CD11b^+^CD11c^+^) from imiquimod-untreated (IMQ–) WT (n=6) and Fcrl5 Tg (n=6) or -treated (IMQ+) WT (n=10) and Fcrl5 Tg (n=7) mice. Data are pooled from two or three independent experiments. (D) Representative flow cytometry plots (left) and absolute numbers (right) of CD4^+^ T cell subsets: effector T (CD4^+^CD25^−^CD62L^−^CD44^hi^), Tfh (CD4^+^CXCR5^+^PD-1^+^), Treg (CD4^+^CD25^+^Foxp3^+^), CD4^+^IFN-γ^+^, and CD4^+^IL-2^+^ cells from imiquimod-untreated (IMQ–) WT (n=6–9) and Fcrl5 Tg (n=6–9) or -treated (IMQ+) WT (n=6–13) and Fcrl5 Tg (n=7–14) mice. Data are pooled from two or three independent experiments. Statistical data are shown as mean values with s.d., and data were analyzed by one-way ANOVA with Tukey’s multiple comparisons test (A-D) or Brown-Forsythe and Welch ANOVA with Dunnett’s T3 multiple comparisons test (C). *P<0.05, **P<0.01, ***P<0.001, ****P<0.0001.

### Impaired B cell anergy to self-antigen in Fcrl5 Tg mice

Anergy is essential for the suppression of self-antigen-recognizing B cells and for limiting autoimmune diseases. To determine whether Fcrl5 overexpression affects B cell anergy, we crossed Fcrl5 Tg mice with MD4 and soluble HEL (ML5) Tg mice. Serum titer analysis showed that MD4 B cells produced HEL-specific IgM antibodies in the absence of the HEL self-antigen (Fig. 5 A); however, the antibody levels decreased in MD4/ML5 double Tg mice, indicating that HEL-specific B cells developed in the presence of the self-antigen becoming anergic and producing fewer HEL-specific antibodies as described previously (Goodnow et al., 1988). We found that crossing MD4/ML5 Tg mice with Fcrl5 Tg mice restored the levels of HEL-specific antibodies (Fig. 5 A). To detect the deletion of peripheral autoreactive B cells, we examined the number of each B cell subset in these mice. In MD4/ML5 Tg mice, the number of total splenic B, FO B, and MZ B cells decreased, whereas in MD4/ML5/Fcrl5 Tg mice, the number was restored, except for MZ B cells (Fig. 5 B). Additionally, the number of CD23^−^CD21^lo^B cells as well as FO B cells increased significantly in MD4/ML5/Fcrl5 Tg mice compared with that in the other mice (Fig. 5 B). These data suggest that Fcrl5 overexpression causes a break in B cell anergy.

**Figure 5.**
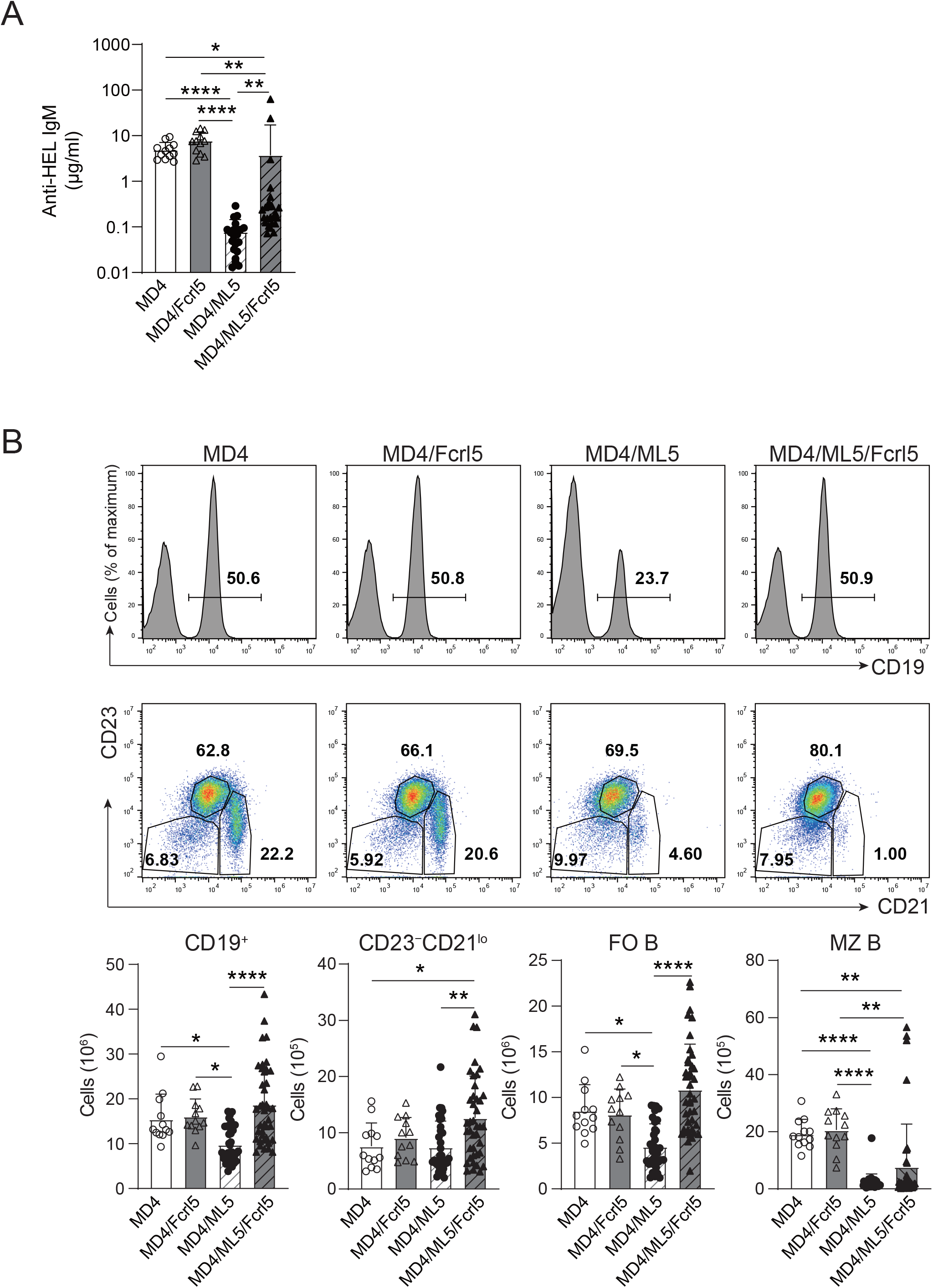
B cell anergy is disrupted in Fcrl5 Tg mice. (A) Anti-HEL antibody production in serum isolated from MD4 Tg (n=12), MD4/Fcrl5 Tg (n=12), MD4/ML5 Tg (n=21), and MD4/ML5/Fcrl5 Tg (n=25) mice, assayed by ELISA. OD, optical density. Data are pooled from six independent experiments. (B) Top, representative flow cytometry plots and absolute numbers of splenic CD19^+^, CD23^−^CD21^lo^ (CD19^+^CD93^−^CD23^−^CD21^lo^), FO B (CD19^+^CD93^−^CD23^+^CD21^lo^), and MZ B (CD19^+^CD93^−^CD23^−^CD21^hi^) cells from MD4 Tg (n=12), MD4/Fcrl5 Tg (n=12), MD4/ML5 Tg (n=35), and MD4/ML5/Fcrl5 Tg (n=40) mice. Data are pooled from eight independent experiments. Statistical data are shown as mean values with s.d., and data were analyzed by Kruskal-Wallis with Dunn’s multiple comparisons test (A) or one-way ANOVA with Tukey’s multiple comparisons test (B). *P<0.05, **P<0.01, ****P<0.0001.

Splenic B cells in MD4/ML5 Tg mice showed typical phenotypes of anergic B cells, as they substantially downregulated cell-surface IgM (Fig. 6 A) and failed to proliferate or upregulate activation marker expression after BCR stimulation (Fig. 6, B and C). Conversely, B cells in MD4/ML5/Fcrl5 Tg mice were not anergic to the HEL antigen, evidenced by higher levels of cell-surface IgM (Fig. 6 A) and self-antigen-mediated proliferation (Fig. 6 B). Additionally, upon HEL stimulation, B cells of MD4/ML5/Fcrl5 Tg mice, but not those of the MD4/ML5 Tg mice, efficiently upregulated CD69, CD86, MHC-II, and PD-L1 and downregulated the inducible costimulatory ligand (ICOSL) (Fig. 6 C). Furthermore, naive OT-II CD4^+^ T cells co-cultured with B cells from the MD4/ML5/Fcrl5 Tg mice differentiated most efficiently into Th1 cells than those from other Tg mice in an ovalbumin concentration-dependent manner (Fig. 6 D). Therefore, Fcrl5 upregulation in B cells reversed the anergic state of autoreactive B cells in the MD4/ML5/Fcrl5 Tg mice, allowing them to produce autoantibodies and potentially present self-antigens that may be involved in autoimmunity.

**Figure 6.**
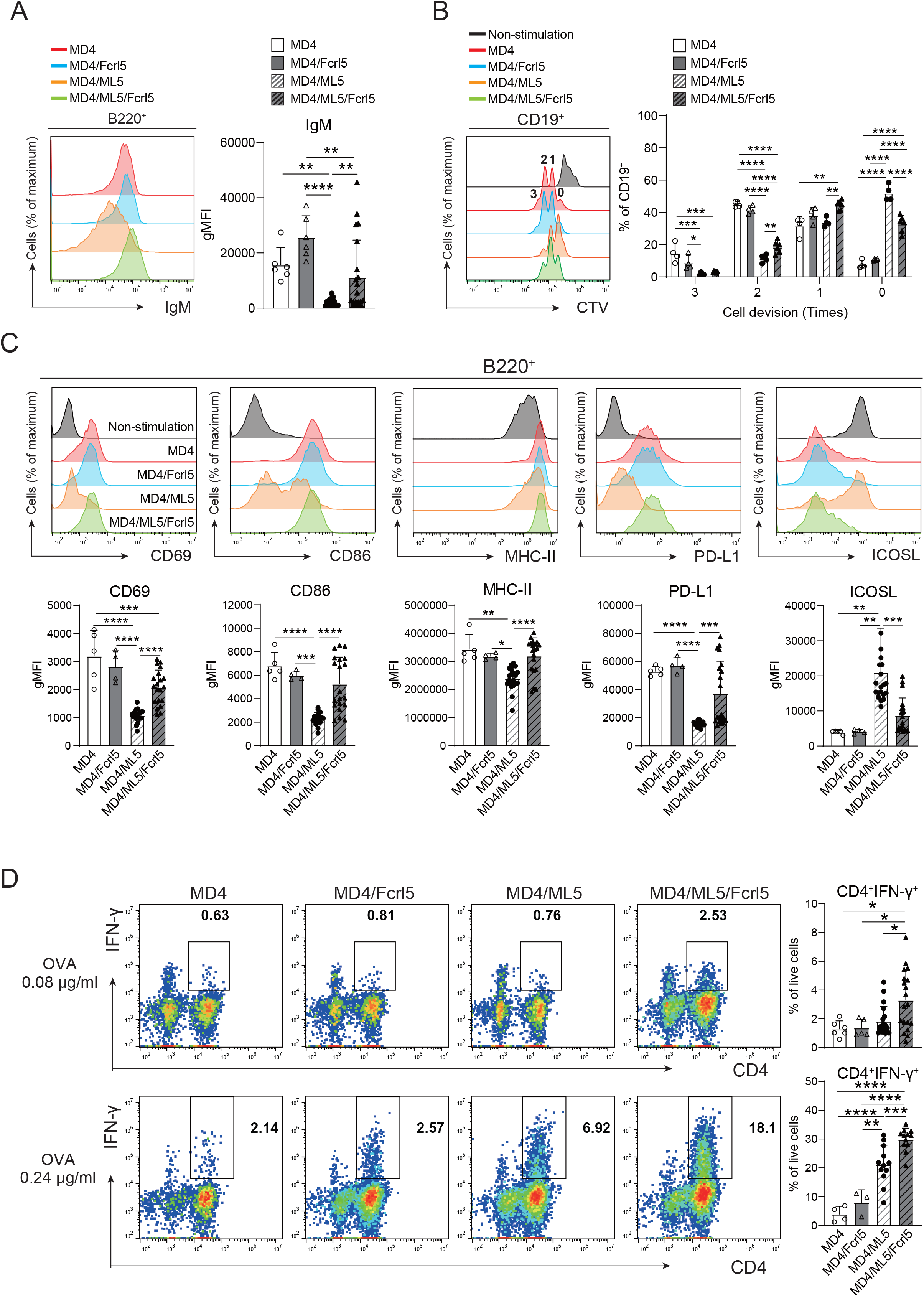
Fcrl5 upregulation causes a break in B cell anergy. (A) A representative histogram and gMFI of cell-surface IgM on B cells from MD4 Tg (n=6), MD4/Fcrl5 Tg (n=6), MD4/ML5 Tg (n=21), and MD4/ML5/Fcrl5 Tg (n=21) mice. gMFI, geometric mean fluorescence intensity. Data are pooled from three independent experiments. (B) Proliferation of CD19^+^ cells isolated from MD4 Tg (n=4), MD4/Fcrl5 Tg (n=4), MD4/ML5 Tg (n=4), and MD4/ML5/Fcrl5 Tg (n=6) mice. CD19^+^ cells are stimulated with HEL + IL-4. Data are pooled from three independent experiments. (C) Representative histograms and gMFIs of CD69, CD86, MHC-Ⅱ, PD-L1, and ICOSL expression level on splenic CD19^+^ cells isolated from MD4 Tg (n=5), MD4/Fcrl5 Tg (n=4), MD4/ML5 Tg (n=20), and MD4/ML5/Fcrl5 Tg (n=20) mice after HEL stimulation. Data are pooled from three independent experiments. (D) Frequencies of CD4^+^IFN-γ^+^ cells derived from co-culture of OT-II T cells in the presence of OVA peptide with CD19^+^ cells purified from MD4 Tg (n=4–6), MD4/Fcrl5 Tg (n=3–5), MD4/ML5 Tg (n=12–19), and MD4/ML5/Fcrl5 Tg (n=12–19) mice. Data are pooled from two or three independent experiments. Statistical data are shown as mean values with s.d., and data were analyzed by one-way ANOVA with Tukey’s multiple comparisons test. *P<0.05, **P<0.01, ***P<0.001, ****P<0.0001.

### Fcrl5 ligation augmented TLR7/9-mediated B cell activation

Given that TLR7/9 activation has been implicated in the development and regulation of autoimmune diseases via anergy disruption (Deane et al., 2007; Rui et al., 2003) and that hFcrl signaling increases TLR-mediated B cell activation (Dement-Brown et al., 2011; Li et al., 2013), we investigated the effect of Fcrl5 on TLR7/9 signaling. As the Fcrl5 ligand was not identified, we stimulated B cells from the Fcrl5 Tg mice with a biotinylated anti-Fcrl5 F(ab’)_2_ antibody cross-linked with streptavidin in the presence of CpG oligodeoxynucleotides (TLR9 ligand) or imiquimod (TLR7 ligand). Co-stimulation with CpG and Fcrl5 resulted in increased cell activation (CD69, CD80, and CD86 expression) and proliferation compared with stimulation with CpG alone (Fig. 7, A and B). TLR7-stimulated CD80 expression, but not CD69 and CD86 expression or proliferation, was promoted by Fcrl5 co-stimulation (Fig. 7, A and B). Hence, Fcrl5 promotes B cell activation in response to TLR signaling, suggesting that Fcrl5 upregulation may contribute to TLR-mediated B cell activation and autoimmunity.

**Figure 7.**
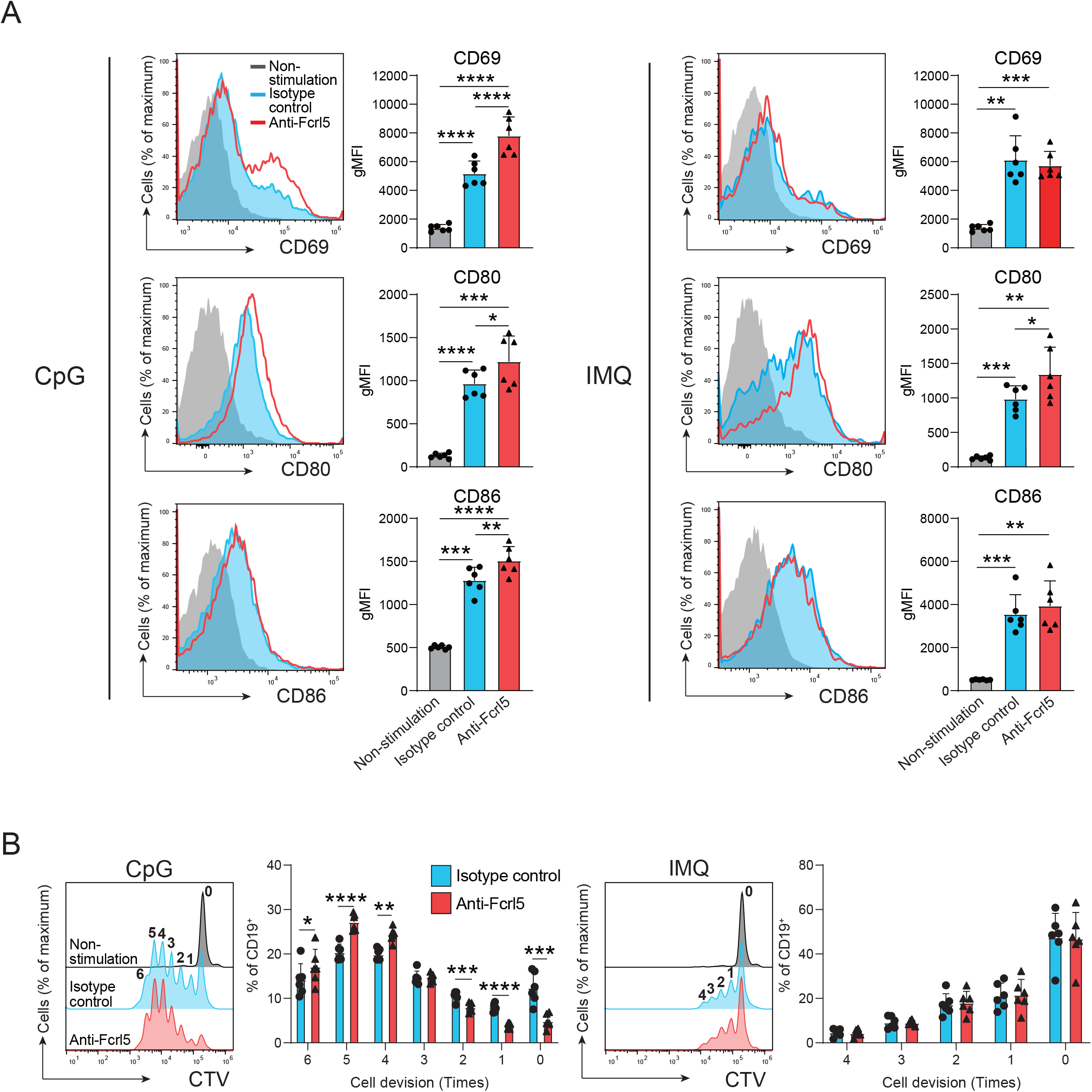
Fcrl5 ligation augments TLR7/9 signaling. (A) Representative histograms and gMFIs of CD69, CD80, and CD86 expression levels on splenic CD19^+^ cells from Fcrl5 Tg mice (n=6) stimulated with CpG (left) or imiquimod (IMQ) (right) in the presence of anti-Fcrl5 F(ab’)_2_ Biotin or isotype control. Data are pooled from two independent experiments. (B) Proliferation of CD19^+^ cells from Fcrl5 Tg mice (n=6) stimulated with CpG (left) or imiquimod (IMQ) (right) in the presence of anti-Fcrl5 F(ab’)_2_ Biotin or isotype control. Data are pooled from two independent experiments. Statistical data are shown as mean values with s.d., and data were analyzed by one-way ANOVA with Tukey’s multiple comparisons test (A) or paired t-test (B). *P<0.05, **P<0.01, ***P<0.001, ****P<0.0001.

## Discussion

Here, we demonstrated that Fcrl5 overexpression in B cells contributed to autoimmune disease development. Our findings that Fcrl5 is more highly expressed in ABCs of aged mice and that B cell-specific Fcrl5 Tg mice show an autoimmune phenotype highlight the importance of the pathogenesis of autoimmune diseases. Additionally, our results showed that Fcrl5 overexpression in B cells can interfere with B cell anergy, suggesting that Fcrl5 plays an important role in regulating B cell-intrinsic peripheral self-tolerance.

Anergy plays a crucial role in maintaining immune tolerance by preventing the production of autoantibodies (Cambier et al., 2007). Breaching this system is a complex and multi-factor process. B cell-mediated autoimmunity is often associated with excessive B cell activation. In this regard, mice deficient in negative regulators of BCRs, such as SHP-1, SHIP-1, PTEN, NFAT1, CD72, CD22, and FcγRIIb, display BCR-stimulated B cell hypersensitivity and develop systemic autoimmune diseases (Cyster and Goodnow, 1995; Getahun et al., 2016; Akerlund et al., 2015; Borde et al., 2006; Li et al., 2008; O’keefe et al., 1999; Bolland and Ravetch, 2000). Mice with mutations that positively regulate BCR signaling—including the ectopic over activation of PI3Kδ and overexpression of CD19—also show aberrant B cell activation, thereby disrupting self-tolerance (Lau et al., 2020; Inaoki et al., 1997). Remarkably, Lyn tyrosine kinase participates in both activation- and inhibition-related signal transductions in B cells (Latour and Veillette, 2001), and both Lyn-deficient and -gain of function mice exhibit disrupted self-tolerance, leading to the manifestation of profound autoimmune disorders (Nishizumi et al., 1995; Hibbs et al., 1995, 2002; Lamagna et al., 2014). This suggests that Lyn is key to maintaining the equilibrium between positive and negative signaling pathways in B cells to control B cell tolerance. Fcrl5 contains ITAM and ITIM sequences in its intracellular region, which are tyrosine phosphorylated upon co-ligation with BCRs to recruit Lyn and SHP-1 (Zhu et al., 2013). Consequently, SHP-1 and Lyn activation mediated by Fcrl5 downregulated BCR signaling in MZ B cells. Conversely, in B1 cells stimulated with BCRs and Fcrl5, SHP-1 inhibited BCR signaling, whereas Lyn promoted it, showing an opposite effect on these cells. This implies that the balance between ITAM-mediated activation signals and ITIM-mediated inhibitory signals by Fcrl5 determines the outcome of B cell activation. Given our finding that Fcrl5 overexpression disrupts immune tolerance, it is conceivable that Fcrl5 in anergic B cells, which are inherently restrained from autoreactive responses, may facilitate BCR signaling and precipitate autoimmune diseases.

A previous RNA-seq analysis revealed that anergic B cells derived from MD4/ML5 Tg mice showed increased *Fcrl5* mRNA levels compared with those shown by naive FO B cells (Hodgson et al., 2022). Nevertheless, our flow cytometric analysis failed to detect the Fcrl5 protein in anergic B cells within the same model (data not shown). This suggests that Fcrl5 does not exhibit its function in typical anergic B cells but that aberrant Fcrl5 upregulation in these cells contributes to their escape from the anergic state. ABCs, presumed pathogenic B cells in autoimmune diseases, show characteristics similar to those of anergic B cells in terms of unresponsiveness to BCRs. The relationship between ABCs and anergic B cells is unknown; however, given that mouse ABCs and human ABC-like cells found in patients with SLE express Fcrl5 and hFcrl3/5, respectively (Wang et al., 2018), it is plausible that Fcrl5-mediated anergy disruption may lead to ABC expansion.

The observations that TLR7-driven autoimmunity was accelerated by B cell-specific Fcrl5 overexpression in mice and that TLR7/9-mediated B cell activation was facilitated by Fcrl5 cross-linking are consistent with many previous findings. This implicates TLR7 and TLR9 signaling in B cells, contributing to autoreactive B cell activation in various lupus mouse models (Deane et al., 2007; Soni et al., 2020). Notably, excessive TLR7 signaling via a gain-of-function mutation in mice causes systemic autoimmune diseases and leads to the B cell autonomous accumulation of ABCs, considered a major source of pathogenicity (Brown et al., 2022). Moreover, TLR7 signaling activation enables B cells to survive by binding to self-antigens via their surface BCRs. Therefore, Fcrl5 may strengthen TLR-mediated signaling and positively regulate BCR signaling to alter the responsiveness threshold to self-antigens. Regarding the action point of Fcrl5, crosslinking it enhances B cell activation when stimulated by TLR stimulation (Fig. 7, A and B) but not by BCR (data not shown) *in vitro*. On the other hand, Fcrl5 overexpression resulted in a break in B cell tolerance in the MD4/ML5 model. The following idea may explain the contradiction at first glance that a signal event formed by the cooperation of TLRs and Fcrl5 might reinforce BCR-mediated activation of anergic B cells upon self-antigen recognition, resulting in a break in tolerance as the anergic B cells were suggested to receive disruptive TLR signaling for the preservation of tolerance (Schwickert et al., 2019). This implies a signaling hierarchy in Fcrl5, TLR, and BCR, although a detailed molecular interplay remains unknown.

Endogenous ligands for mouse Fcrl5 have not been identified; however, hFcrl3 and hFcrl5 have been reported to bind to secretory IgA and IgG, respectively (Agarwal et al., 2020; Wilson et al., 2012; Franco et al., 2013). When BCR and hFcrl3/5 are co-stimulated, hFcrl3/5 can inhibit BCR signaling via their ITIMs (Kochi et al., 2009; Haga et al., 2007). Conversely, hFcrl3/5 promote innate TLR9 signaling (Li et al., 2013; Dement-Brown et al., 2011). The simultaneous stimulation of hFcrl3 and TLR9 activates B cells via the NF-κB and mitogen-activated protein kinase pathways. (Li et al., 2013). Furthermore, the cross-linking of hFcrl5 and BCRs in TLR9-stimulated B cells increases B cell proliferation. These data suggest that hFcrl3/5 modulate TLR9 signaling. Additionally, hFcrl5 constitutively associates with the activating BCR co-receptor CD21 and functions together with CD21 to promote BCR signaling (Franco et al., 2018). In this context, mouse Fcrl5 may cooperate with CD21 to promote BCR signaling to breach anergy *in vivo*.

Our study provides important insights into the role of Fcrl5 in breaking B cell anergy and its effect on the pathogenesis of autoimmune diseases. The mechanism of action of Fcrl5 in activating autoreactive B cells is unclear; nonetheless, Fcrl5 can modulate BCR and TLR signaling. Elucidation of the endogenous ligands for Fcrl5 will be important in understanding its role in these signaling pathways.

## Acknowledgments

We thank Tanaka, M. and Kageyama, K. for Technical Support in the Medical Institute of Bioregulation, Kyushu University for the animal facility: Furuno, N. for secretarial help. We acknowledge the NGS core facility at the Research Institute for Microbial Diseases of Osaka University for the sequencing and data analysis. This work was partially supported by JSPS KAKENHI (JP18H02626, 19K22537, JP21H02753 to Y.B., JP22H03112, JP22K19547 to S.T. and 18790697 to Y.K.), JSPS Fellows (JP21J20013 and JP22KJ2376 to C.O), JST FOREST Program (JPMJFR210S to S.T.), Agency for Medical Research and Development (AMED) (JP19ek0410044 and JP19gm6110004 to Y.B.), the Medical Research Center Initiative for High Depth Omics and RIIT (to Y.K and Y.B.), Kato Memorial Bioscience Foundation, and The Naito Foundation (to Y.B.).

## Author contributions

C.O. performed experiments, analyzed data, and wrote the manuscript. T.S. designed the study, supported experiments, and provided important inputs. K.M. support the experiments and screened anti-Fcrl5 antibody. T.I. and Y.O performed H&E staining and histopathological diagnosis. M.O analyzed ImmuNextUT data. K.Y. designed the study and provided important inputs. Y.K. designed the study, performed experiments, and analyzed data. Y.B. designed and supervised the study and wrote the manuscript. All authors reviewed and commented upon the manuscript.

## Disclosures

The authors declare that they have no competing interests.

## Material and methods

### Mice

C57BL/6 wild-type (WT) mice were obtained from CLEA Japan, Inc. Fcrl5 transgenic (Tg) mice were generated by using B-cell specific expression vector, pEµIgH containing the human Eµ enhancer (IgH intronic enhancer) and mouse IgVH promoter as previously described (Koike et al., 1995). Briefly, murine Fcrl5 cDNA was subcloned into the EcoRI-digested DNA fragment at the EcoRI site of pEµIgH. Then, the linearized pEµIgH/Fcrl5 DNA fragments were injected into fertilized eggs of C57BL/6 mice. The offspring mice were screened by PCR analyses using tail DNA and B cell-specific Fcrl5 expression was confirmed by flow cytometric analysis. MD4 Tg mice were crossed with soluble HEL (ML5) Tg mice to obtain MD4/ML5 Tg mice (Goodnow et al., 1988). MD4 Tg and MD4/ML5 Tg mice were subsequently bred with Fcrl5 Tg mice to obtain MD4/Fcrl5 Tg and MD4/ML5/Fcrl5 Tg mice. OT-II TCR-Tg mice were described previously (Barnden et al., 1998). 50–100-week-old WT and Fcrl5 Tg mice were used for analysis of gene expression, B cell development, Ig production and pathological analysis of aged mice, shown in Figs. 1, 2, S1 and S3. 8–24-week-old MD4 Tg, MD4/Fcrl5 Tg, MD4/ML5 Tg and MD4/ML5/Fcrl5 Tg mice were used to examine the role of Fcrl5 in B cell anergy, shown in Figs. 5 and 6. Other mice were used for experiments at 8–12-week-old. All mice used in the experiments were kept under specific pathogen– free conditions. All experiments were conducted in accordance with the animal experiment guidelines with the approval of the Animal Ethics Committee of Kyushu University School of Medicine.

### *In vivo* treatment with TLR7 agonist Imiquimod

WT, Fcrl5 Tg, and MD4 Tg mice were treated with 1.25 mg of 5% imiquimod cream (Mochida Pharmaceuticals, Tokyo, Japan) on right ear once every 2 days for 4 or 10 weeks.

### Cell staining and cell sorting

Single-cell suspensions were prepared and resuspended in FACS staining buffer. Cells were incubated with optimal concentrations of antibodies to the following targets: B220 (RA3-6B2), CD4 (GK1.5), CD5 (53-7.3), CD8a (53-6.7), CD11b (M1/70), CD11c (N418), CD19 (6D5 or 1D3), CD21/35 (7E9), CD23 (B3B4), CD25 (PC61), CD43 (eBioR2/60 or S7), CD44 (IM7), CD62L (MEL-14), CD69 (H1.2F3), CD80 (16-10AJ), CD86 (GL-1), CD93 (AA4.1), CD138 (281-2), CXCR5 (2G8), FAS (SA367H8), Foxp3 (FJK-16S), GL7 (GL7), I-A/I-E (M5/114.15.2), ICOSL (HK5.3), IFN-γ (XMG1.2), IgG (Poly4053), IgM (RMM-1), IL-2 (JES6-5H4), NK1.1 (PK136), PD-1 (29F.1A12), PD-L1 (10F.9G2), TACI (eBioBF10-3), TER-119 (TER-119) (all from BioLegend, BD Biosciences, eBioscience and Invitrogen) and Fcrl5 (128). IgG2a isotype control (RTK2758) and Rat IgG F(ab’)_2_ Biotin isotype control (Rockland) were used as a control of Fcrl5 staining. B cells were enriched with CD19 Microbeads (Miltenyi Biotec) or MojoSort Mouse Pan B cell Isolation Kit (BioLegend), following the manufacturer’s instructions. FO B (CD19^+^B220^hi^CD43^−^ CD23^+^CD21^lo^) cells and ABCs (CD19^+^B220^hi^CD43^−^CD23^−^CD21^lo^CD11b^+^CD11c^+^) were purified by FACS Melody (BD Biosciences) as shown in Fig. S1 A.

### Production of monoclonal antibodies to Fcrl5

The full-length coding sequence of Fcrl5 was cloned into pDisplay vector (Thermo Fisher Scientific) and then transfected into HEK-293 cells (JCRB Cell Bank) to establish a stable transfectant line (293-Fcrl5 cells). Wistar rats were immunized with the 293-Fcrl5 cells, and hybridomas were generated by fusing splenic B-cells with mouse X-63 myeloma cells. The supernatants of each hybridoma were screened by ELISA using 293-Fcrl5 cells as antigens and further evaluated by flow cytometric analysis using a stable transfectant line expressing Fcrl5 in ST486 cells (ATCC). The antibody specificity was also evaluated using stable transfectants of Fcrl1 and Fcrl2 in HEK-293 cells, and the clone 128 was selected. The 128 hybridoma cells were injected into the peritoneal cavity of Pristan-primed BALB/c mice, and antibodies in the ascitic fluid were purified by protein A column chromatography. F(ab’)_2_ fragment antibodies were generated by pepsin digestion of whole IgG antibodies. All the work from rat immunization to antibody fragmentation was conducted at Immuno-Biological Laboratories Co, Ltd. These antibodies were biotinylated using the EZ-Link Sulfo-NHS-LC-Biotin Kit (Thermo Fisher Scientific).

### In vitro B cell culture and stimulation

The isolated B cells were cultured in 10% FCS RPMI [1 mM sodium pyruvate solution (Nacalai Tesque) penicillin-streptomycin mixed solution (Nacalai Tesque), glutamine, 10 mM HEPES (Nacalai Tesque), 1 × MEM essential amino acid solution (Nacalai Tesque), 50 μM 2-Mercaptoethanol (Sigma-Aldrich). For B cell activation assay, B cells isolated from MD4 Tg, MD4/Fcrl5 Tg, MD4/ML5 Tg, and MD4/ML5/Fcrl5 Tg mice were stimulated with 0.1 μg/ml HEL (Sigma-Aldrich) for 15-18 h to detect CD69, CD80, CD86, MHC-Ⅱ, PD-L1 and ICOSL expression. For proliferation assay, B cells were labeled with 1μM Cell Trace Violet (Invitrogen) and stimulated with 0.3 μg/ml HEL plus 20 ng/ml IL-4 (R&D Systems), 0.2 μg/ml CpG (InvivoGen) or 1 μg/ml Imiquimod (IMQ) (InvivoGen) in the presence of 22.5 μg/ml anti-Fcrl5 F(ab’)_2_ Biotin or Rat IgG F(ab’)_2_ Biotin isotype control (Rockland) and 20 μg/ml streptavidin (Thermo Fisher Scientific) for 72 h.

### Naive CD4 ^+^ T cell isolation and in vitro B-T cell coculture

Splenic CD4^+^ T cells derived from OT-II TCR-Tg mice were enriched with Biotin-labeled antibody cocktail (B220, CD8a, CD11b, CD11c, CD19, CD25, NK1.1, and TER-119) with Streptavidin Microbeads (Miltenyi Biotec). CD4^+^ T cells were further incubated with CD62L Microbeads (Miltenyi Biotec) to enrich CD62L^+^CD4^+^ T cells. These CD62L^+^CD4^+^ cells were labeled with 1μM Cell Trace Violet (Invitrogen) and co-cultured with B cells in the presence of 0.08 and 0.24 μg/ml OVA_323–339_-peptide (MBL) for 3 days.

### Intracellular staining in T cells

Cells were stimulated with 500 ng/ml ionomycin and 50 ng/ml PMA in the presence of monensin (eBioscience) for 4 h. Cells were then permeabilized with IC fixation buffer/permeabilization buffer (eBioscience) and analyzed for the expression of intracellular cytokines with anti-IFN-γ or anti-IL-2 antibody. Intracellular transcription factors were detected using the Foxp3 staining buffer kit with anti-Foxp3 antibody.

### Histological analysis

Tissue specimens were fixed in 10% neutral buffered formalin and embedded in paraffin. Tissue sections were stained with hematoxylin–eosin. Stained slide images were obtained using Leica Aperio GT450 (Leica, Tokyo, Japan) slide scanner at 40× objective magnification, an Olympus DP27 camera (Evident, Tokyo, Japan), or an all-in-one fluorescence microscope (BZ-X800, KEYENCE, Osaka, Japan). For liver analysis, the largest and second-largest cut surfaces, when cut at the long axis of the liver, were used for measurement. For kidney analysis, the cut surface, when cut at the minor axis of the bilateral kidneys, was used for measurement. The total area and the area of lymphoid cell infiltration were measured using the image analysis software QuPath (Bankhead et al., 2017). Lung tissue distortion was assessed by measuring the mean linear intercept between the alveolar walls (Lm) at 20× magnification. For each sample, intercepts sampled in four views were measured using NIS Element (Nikon, Tokyo, Japan). For lymphoid cell aggregated area analysis of the lung, each figure for analysis was captured at 10× magnification, and the area was measured using NIS Element (Nikon, Tokyo, Japan).

### Detection of autoantibodies with human epithelial type 2 (HEp-2) slides

Diluted mouse serum (1:100 in 0.1% BSA in PBS) was incubated on HEp-2 slides (MBL or BioRad) for 1 hour in the dark at room temperature. Subsequently, the slides were rinsed once with a squirt bottle and washed thrice for 10 min with PBS. For the detection of mouse IgG, the slides were incubated for 1 hour in the dark at room temperature with an Alexa 488-conjugated goat anti-mouse IgG antibody (BioLegend). Following three washing steps, mounting medium (MBL or BioRad) was added, together with a cover slip. Images were acquired with an all-in-one fluorescence microscope (BZ-X800, KEYENCE, Osaka, Japan). Images were evaluated as negative (Negative) or positive (Positive) reactive to antigen. Positive images were classified as reactive to cytoplasmic antigen (cytoplasmic), reactive to nuclear and cytoplasmic antigen (nuclear and cytoplasmic), or reactive to nuclear and/or nucleolar/nuclear antigen (nucleolar/nuclear) (Domeier et al., 2018).

### ELISA

Plates were coated with 10 μg/ml dsDNA (Invitrogen), 2 μg/ml Histon (Roche), 1 μg/ml Sm/RNP (GenWay), 10 μg/ml HEL or 0.5 μg/ml anti-mouse IgM, IgG1, IgG2b, IgG2c, IgG3, IgA (SouthernBiotech) in PBS at 4℃ overnight. The serum diluted and incubated on coated plates at 4℃ overnight. Plates were then incubated for 3 h with horseradish peroxidase–labeled goat anti-mouse IgM, IgG, IgG1, IgG2b, IgG2c, IgG3, or IgA (SouthernBiotech). OD_450_ was measured on a microplate reader (iMArkTM Microplate Reader, Bio-Rad).

### qPCR and RNA-seq

Total RNA was isolated from cells using miRNeasy Micro Kit (Qiagen) according to the manufacturer’s protocol. Full-length cDNAs were prepared using the SMART-Seq HT Kit (Takara Bio, Mountain View, CA) according to the manufacturer’s instructions. Real-Time PCR was performed using the THUNDERBIRD^TM^ SYBR^®^ qPCR MIX (TOYOBO). The following custom primers were used: *Fcrl1* forward (5’-CACACGGAGTAAGTGAGTCCT-3’), *Fcrl1* reverse (5’-TCAGGCCTTGGGCTTGTATG-3’); *Fcrl5* forward (5’-GCATGCACGTGAACGTAGAG-3’), *Fcrl5* reverse (5’-TCCTTCAAACACAGCAGGTG-3’); *Fcrla* forward, (5’-GAAGCAGAGCCCACAACTG-3’), *Fcrla* reverse (5’-GTGGGAGGAGTTTCCGAAG-3’). For global gene expression analysis, an Illumina library was prepared using a Nextera DNA Library Preparation Kit (Illumina, SanDiego, CA) according to SMARTer kit instructions. Sequencing was performed on an Illumina NovaSeq 6000 sequencer (Illumina) in the 101-base single-end mode. Sequenced reads were mapped to the mouse reference genome sequences (mm10) using TopHat v2.0.13 in combination with Bowtie2 ver. 2.2.3 and SAMtools ver. 0.1.19. The fragments per kilobase of exon per million mapped fragments (FPKMs) were calculated using Cufflinks version 2.2.1.

### Regulatory effect of GWAS SNP in immune cells

The regulatory effect (eQTL effect) of GWAS SNP rs7528684 and rs11264750 in the promoter region of *hFcrl3* and *hFcrl5* were evaluated by using the dataset of ImmunNexUT (Ota et al., 2021) which consists of 19 distinct immune cell subsets from 337 patients diagnosed with 10 immune-mediated diseases and 79 healthy volunteers. The plots were made by the web tools in the ImmuNexUT portal (https://www.immunexut.org/)

### Statistical analysis

P values were calculated with paired or unpaired Student’s t-test or a non-parametric Mann-Whitney test for two-group comparisons and with by one-way ANOVA with Tukey’s multiple comparisons test, Brown-Forsythe and Welch ANOVA with Dunnett’s T3 multiple comparisons test or Kruskal-Wallis with Dunn’s multiple comparisons test for multi-group comparisons. P values of < 0.05 were considered significant; and the following values were delineated: *P < 0.05, **P < 0.01, ***P < 0.001 and ****P < 0.0001. Statistical analysis was performed in the GraphPad Prism software.

## Notes

### Competing Interest Statement

The authors have declared no competing interest.

