## Supplemental Figures for "Upregulated Fcrl5 disrupts B cell anergy and causes autoimmunity"

Figure S1

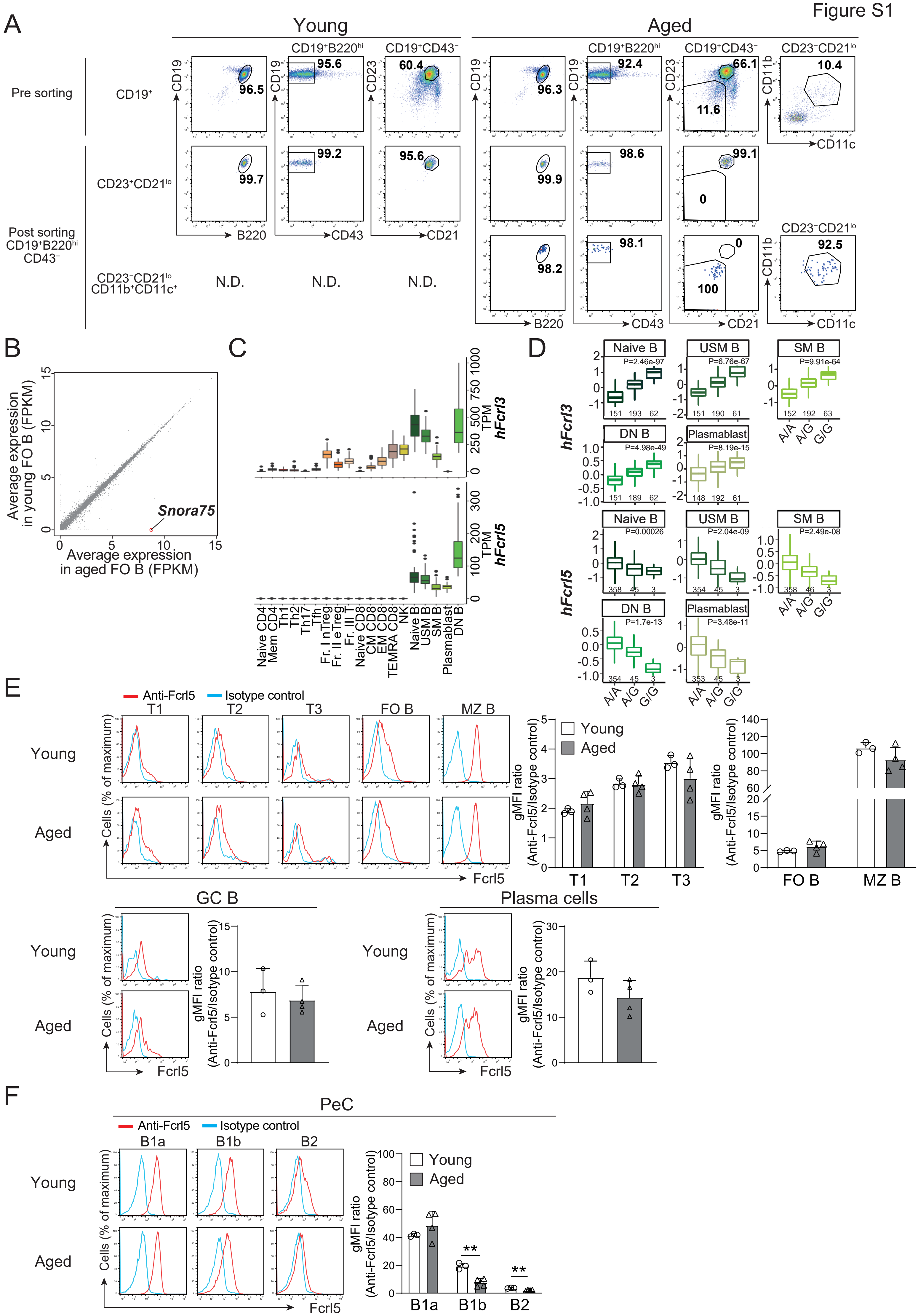
A
Young
Aged
Pre sorting
CD19⁺
CD19⁺B220hi
CD19⁺CD43⁻
CD23⁻CD21lo
Post sorting CD19⁺B220hi CD43⁻
CD23⁺CD21lo
CD23⁻CD21lo CD11b⁺CD11c⁺
N.D.
B220
CD43
CD21
CD11c
CD19
CD23
CD11b
96.5
95.6
60.4
96.3
92.4
66.1
11.6
10.4
99.7
99.2
95.6
99.9
98.6
99.1
0
98.2
98.1
100
92.5
B
Average expression in young FO B (FPKM)
Average expression in aged FO B (FPKM)
Snora75
C
hFcrl3
hFcrl5
TPM
Naive CD4
Mem CD4
Th1
Th2
Th17
Tfh
Fr. I nTreg
Fr. II eTreg
Fr. III T
Naive CD8
CM CD8
EM CD8
TEMRA CD8
NK
Naive B
USM B
SM B
Plasmablast
DN B
D
Naive B
USM B
SM B
DN B
Plasmablast
P=2.46e-97
P=6.76e-67
P=9.91e-64
P=4.98e-49
P=8.19e-15
P=0.00026
P=2.04e-09
P=2.49e-08
P=1.7e-13
P=3.48e-11
A/A
A/G
G/G
E
Anti-Fcrl5
Isotype control
T1
T2
T3
FO B
MZ B
Cells (% of maximum)
Fcrl5
gMFI ratio (Anti-Fcrl5/Isotype control)
GC B
Plasma cells
F
PeC
B1a
B1b
B2
**

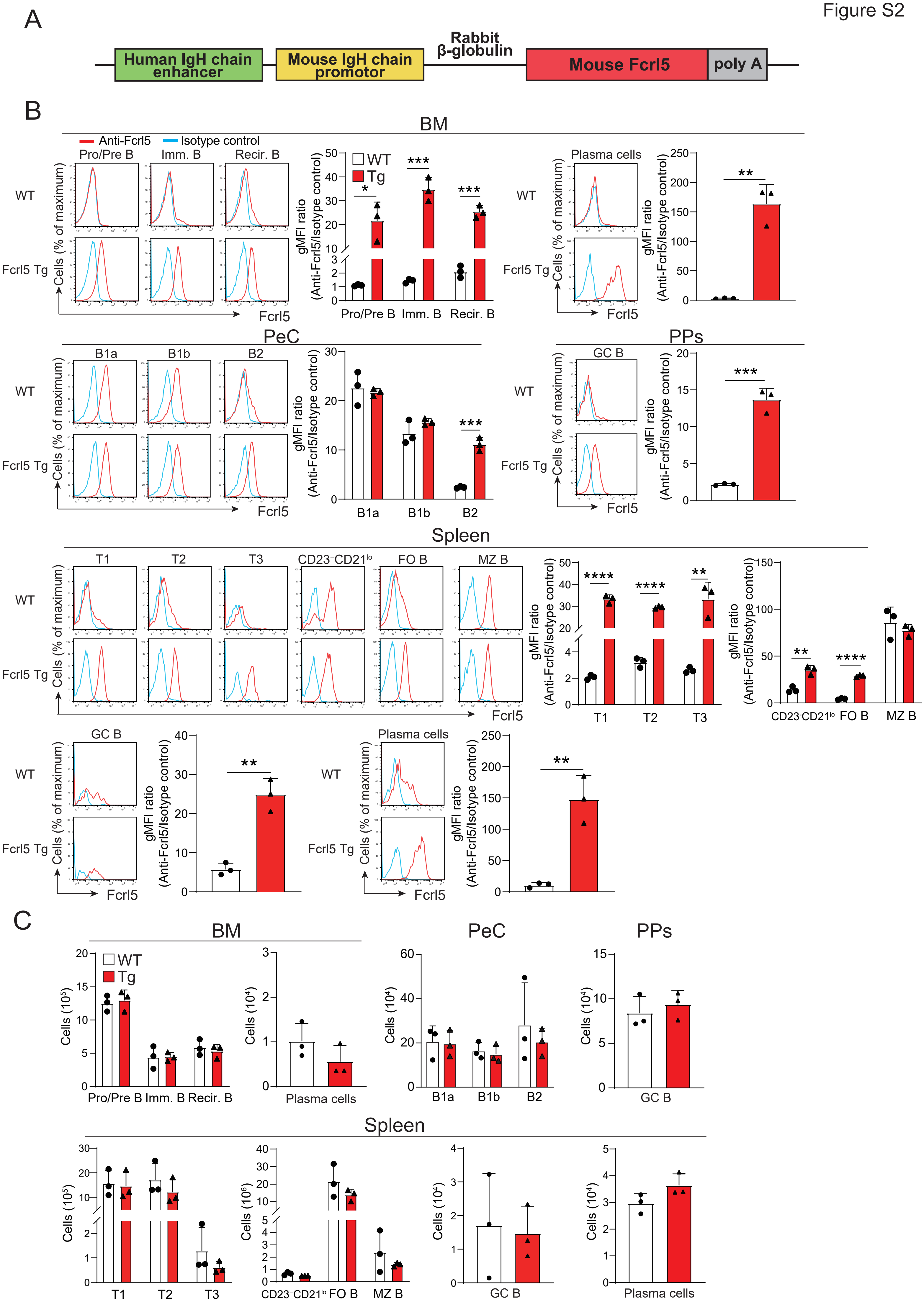
A
Human IgH chain enhancer
Mouse IgH chain promotor
Rabbit β-globulin
Mouse Fcrl5
poly A
B
BM
Anti-Fcrl5
Isotype control
Pro/Pre B
Imm. B
Recir. B
WT
Fcrl5 Tg
Cells (% of maximum)
Fcrl5
gMFI ratio (Anti-Fcrl5/Isotype control)
Tg
Plasma cells
PeC
B1a
B1b
B2
PPs
GC B
Spleen
T1
T2
T3
CD23⁻CD21lo
FO B
MZ B
C
Cells (10⁵)
Cells (10⁴)
Cells (10⁶)

Figure S3

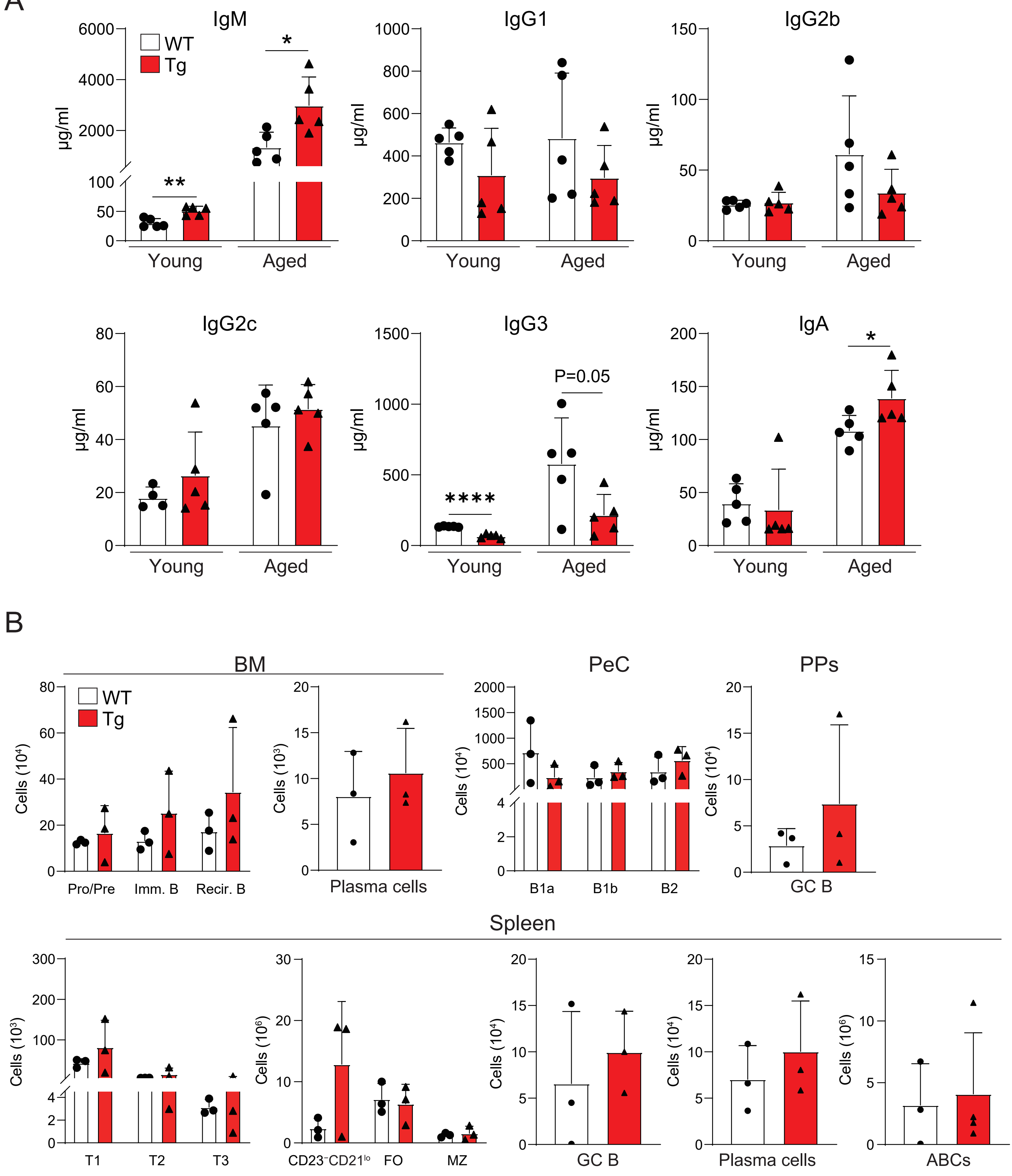
A
IgM
WT
Tg
6000
4000
2000
100
50
0
μg/ml
**
*
Young
Aged
IgG1
1000
800
600
400
200
0
IgG2b
150
100
50
IgG2c
80
60
40
20
IgG3
1500
1000
500
****
P=0.05
IgA
200
150
B
BM
PeC
PPs
Cells (10⁴)
Cells (10³)
Pro/Pre
Imm. B
Recir. B
Plasma cells
2000
1500
1000
500
B1a
B1b
B2
GC B
Spleen
300
T1
T2
T3
Cells (10⁶)
CD23⁻CD21ˡᵒ
FO
MZ
ABCs

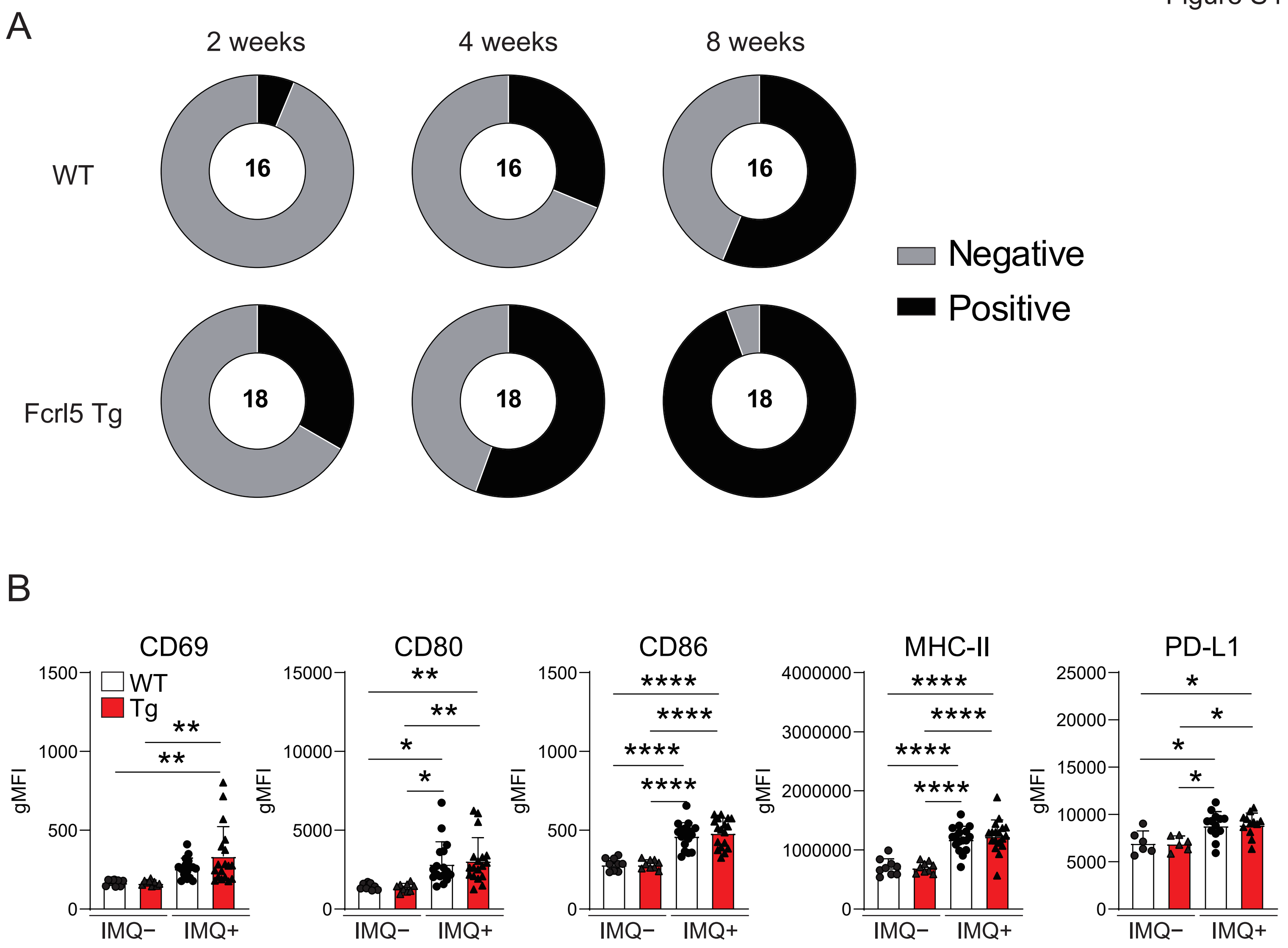
A
2 weeks
4 weeks
8 weeks
WT
16
16
16
Fcrl5 Tg
18
18
18
Negative
Positive
B
CD69
CD80
CD86
MHC-II
PD-L1
WT
Tg
gMFI
IMQ−
IMQ+

Figure S5

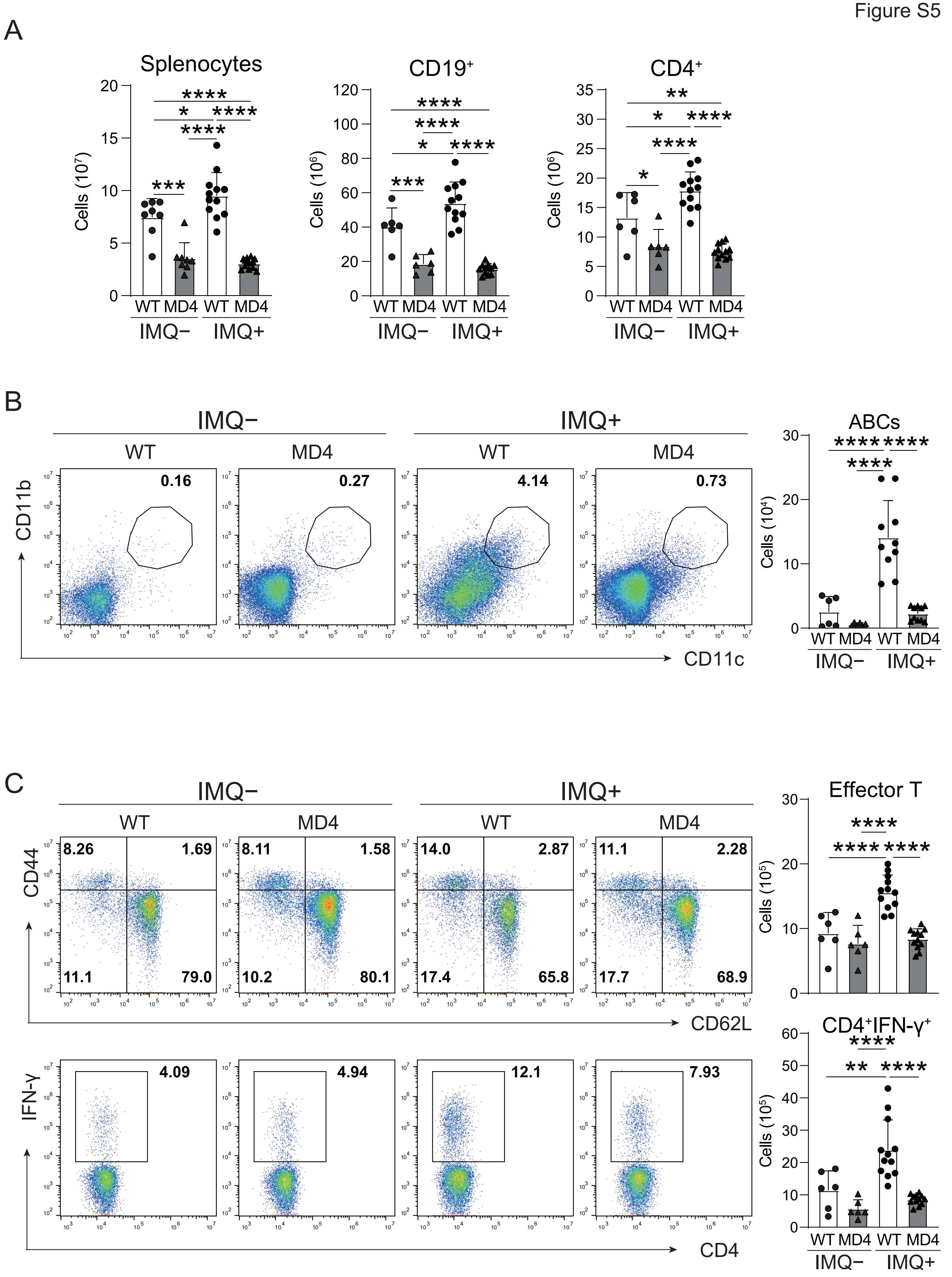
A
Splenocytes
CD19+
CD4+
Cells (10^7)
Cells (10^6)
WT MD4 WT MD4
IMQ− IMQ+
B
IMQ−
IMQ+
WT
MD4
0.16
0.27
4.14
0.73
CD11b
CD11c
ABCs
Cells (10^4)
C
Effector T
CD4+IFN-γ+
Cells (10^5)
CD44
CD62L
8.26
1.69
11.1
79.0
8.11
1.58
10.2
80.1
14.0
2.87
17.4
65.8
11.1
2.28
17.7
68.9
IFN-γ
CD4
4.09
4.94
12.1
7.93

**Supplemental figure legends**

**Figure S1. Gating strategy and distribution of Fcrl5 expression.**

(A) Gating strategy employed to sort FO B (CD19$^{+}$B220$^{hi}$CD43$^{-}$CD23$^{+}$CD21$^{lo}$) cells and ABCs (CD19$^{+}$B220$^{hi}$CD43$^{-}$CD23$^{-}$CD21$^{lo}$CD11b$^{+}$CD11c$^{+}$) from young and aged WT mice (n=3 mice each). (B) Differential gene expression of FO B cells from young WT mice and FO B cells from aged WT mice (n=3 mice each) presented as a scatter plot. (C) Expression of *hFcrl3* and *hFcrl5* in immune subsets: unswitched memory B (USM B), switched memory B (SM B), and double negative B (DN B) cells in healthy individuals (n=79). TPM, Transcript per million. (D) eQTL effect of GWAS SNP rs7528684 (*hFcrl3*) and rs11264750 (*hFcrl5*) in the immune subsets from patients with 10 immunological disorders (n=416). P-values were obtained by testing the alternative hypothesis that the slope of a linear regression model between genotype and expression deviates from 0. (E) Representative histograms and gMFIs of Fcrl5 expression on T1 (CD19$^{+}$CD93$^{+}$IgM$^{+}$CD23$^{-}$), T2 (CD19$^{+}$CD93$^{+}$IgM$^{+}$CD23$^{+}$), T3 (CD19$^{+}$CD93$^{+}$IgM$^{-}$CD23$^{+}$), FO B (CD19$^{+}$CD93$^{-}$CD23$^{+}$CD21$^{lo}$), MZ B (CD19$^{+}$CD93$^{-}$CD23$^{-}$CD21$^{hi}$), GC B (B220$^{+}$Fas$^{+}$GL7$^{+}$), and plasma (CD138$^{+}$TACI$^{+}$) cells from young WT (n=3) and aged WT mice (n=4). Data are representative of two independent experiments. (F) Representative histograms and gMFIs of Fcrl5 expression of B1a (CD19$^{+}$CD5$^{+}$CD23$^{-}$), B1b (CD19$^{+}$CD5$^{-}$CD23$^{-}$), and B2 (CD19$^{+}$CD5$^{-}$CD23$^{+}$) cells in PeC from young WT (n=3) and aged WT (n=4) mice. Data are representative of two independent experiments. Statistical data are shown as mean values with s.d., and data were analyzed by unpaired Student's t-test. **P<0.01.

**Figure S2. Analysis of Fcrl5 expression and B cell development in young Fcrl5 Tg mice.**

(A) A scheme of the constructs used to generate Fcrl5 Tg mice. (B) Representative histograms and gMFIs of Fcrl5 expression on Pro/Pre B (B220$^{+}$IgM$^{-}$), Immature B (Imm. B; B220$^{lo}$IgM$^{+}$), recirculating B (Recir. B; B220$^{hi}$IgM$^{+}$) and plasma cells (CD138$^{+}$TACI$^{+}$) in BM, B1a

(CD19$^{+}$CD5$^{+}$CD23$^{-}$), B1b (CD19$^{+}$CD5$^{-}$CD23$^{-}$), and B2 (CD19$^{+}$CD5$^{-}$CD23$^{+}$) cells in PeC, GC B (B220$^{+}$Fas$^{+}$GL7$^{+}$) in PPs and T1 (CD19$^{+}$CD93$^{+}$IgM$^{+}$CD23$^{-}$), T2 (CD19$^{+}$CD93$^{+}$IgM$^{+}$CD23$^{+}$), T3 (CD19$^{+}$CD93$^{+}$IgM$^{-}$CD23$^{+}$), CD23$^{-}$CD21$^{lo}$ (CD19$^{+}$CD93$^{-}$CD23$^{-}$CD21$^{lo}$), FO B (CD19$^{+}$CD93$^{-}$CD23$^{+}$CD21$^{lo}$), MZ B (CD19$^{+}$CD93$^{-}$CD23$^{-}$CD21$^{hi}$), GC B (B220$^{+}$Fas$^{+}$GL7$^{+}$), and plasma (CD138$^{+}$TACI$^{+}$) cells in spleen from young WT and young Fcrl5 Tg mice (n=3 mice each). Data are representative of four independent experiments. (C) Absolute numbers of B cell subsets described in (B) from young WT and young Fcrl5 Tg mice (n=3 mice each). Data are representative of five independent experiments. Statistical data are shown as mean values with s.d., and data were analyzed by unpaired Student's t-test. *P<0.05, **P<0.01, ***P<0.001, ****P<0.0001.

**Figure S3. Analysis of natural antibody titers in young and aged Fcrl5 Tg mice and B cell development in aged Fcrl5 Tg mice.**

(A) Antibody production in serum isolated from young WT, young Fcrl5 Tg, aged WT, and aged Fcrl5 Tg mice (n=5 mice each), assayed by ELISA. Data are pooled from two independent experiments. (B) Absolute numbers of Pro/Pre B (B220$^{+}$IgM$^{-}$), Immature B (Imm. B; B220$^{lo}$IgM$^{+}$), recirculating B (Recir. B; B220$^{hi}$IgM$^{+}$) and plasma cells (CD138$^{+}$TACI$^{+}$) in BM, B1a (CD19$^{+}$CD5$^{+}$CD23$^{-}$), B1b (CD19$^{+}$CD5$^{-}$CD23$^{-}$) and B2 (CD19$^{+}$CD5$^{-}$CD23$^{+}$) cells in PeC, GC B (B220$^{+}$Fas$^{+}$GL7$^{+}$) in PPs and T1 (CD19$^{+}$CD93$^{+}$IgM$^{+}$CD23$^{-}$), T2 (CD19$^{+}$CD93$^{+}$IgM$^{+}$CD23$^{+}$), T3 (CD19$^{+}$CD93$^{+}$IgM$^{-}$CD23$^{+}$), CD23$^{-}$CD21$^{lo}$ (CD19$^{+}$CD93$^{-}$CD23$^{-}$CD21$^{lo}$), FO B (CD19$^{+}$CD93$^{-}$CD23$^{+}$CD21$^{lo}$), MZ B (CD19$^{+}$CD93$^{-}$CD23$^{-}$CD21$^{hi}$), GC B (B220$^{+}$Fas$^{+}$GL7$^{+}$), plasma (CD138$^{+}$TACI$^{+}$) cells, and ABCs (CD19$^{+}$B220$^{hi}$CD23$^{-}$CD21$^{lo}$CD11b$^{+}$CD11c$^{+}$) in spleen from aged WT (n=3) and aged Fcrl5 Tg (n=3–4) mice. Data are representative of two or three independent experiments. Statistical data are shown as mean values with s.d., and data were analyzed by Mann-Whitney test. *P<0.05, **P<0.01, ****P<0.0001.

**Figure S4. Autoantibody production and B cell activation in imiquimod-treated Fcrl5 Tg mice.** (A) ANAs production in serum from imiquimod-treated (IMQ+) WT (n=16) and Fcrl5 Tg (n=18) for 2, 4, and 8 weeks, detected by immunofluorescence assay using HEp-2 cells. Data are pooled from three independent experiments. (B) Representative histograms and gMFIs of CD69, CD80, CD86, MHC-II, and PD-L1 expression levels on splenic CD19$^{+}$ cells isolated from imiquimod-untreated (IMQ–) WT (n=9) and Fcrl5 Tg (n=9) or -treated (IMQ+) WT (n=17) and Fcrl5 Tg (n=18) mice. Data are pooled from three independent experiments. Statistical data are shown as mean values with s.d., and data were analyzed by one-way ANOVA with Tukey's multiple comparisons test. *P<0.05, **P<0.01, ****P<0.0001.

**Figure S5. Requirement of antigen recognition via BCR for imiquimod-induced inflammation.** (A) Absolute number of splenocytes, CD19$^{+}$, and CD4$^{+}$ cells in spleen from imiquimod-untreated (IMQ–) WT (n=6–8) and MD4 Tg (n=6–8) or -treated (IMQ+) WT (n=12) and MD4 Tg (n=12) mice. Data are pooled from two independent experiments. (B) Representative flow cytometry plots and absolute numbers of ABCs (CD19$^{+}$B220$^{hi}$CD23$^{-}$CD21$^{lo}$CD11b$^{+}$CD11c$^{+}$) from imiquimod-untreated (IMQ–) WT (n=6) and MD4 Tg (n=6) or -treated (IMQ+) WT (n=10) and MD4 Tg (n=10) mice. Data are pooled from two independent experiments. (C) Representative flow cytometry plots (left) and absolute numbers (right) of CD4$^{+}$ T cell subsets: effector T (CD4$^{+}$CD25$^{-}$CD62L$^{-}$CD44$^{hi}$) and CD4$^{+}$IFN-$\gamma^{+}$ cells from imiquimod-untreated (IMQ–) WT (n=6) and MD4 Tg (n=6) or -treated (IMQ+) WT (n=12) and MD4 Tg (n=12) mice. Data are pooled from two independent experiments. Statistical data are shown as mean values with s.d., and data were analyzed by one-way ANOVA with Tukey's multiple comparisons test. *P<0.05, **P<0.01, ***P<0.001, ****P<0.0001.
